# *In situ* architecture of human kinetochore-microtubule interface visualized by cryo-electron tomography

**DOI:** 10.1101/2022.02.17.480955

**Authors:** Wei Zhao, Grant J. Jensen

## Abstract

Faithful segregation of chromosomes during mitosis relies on a carefully coordinated and intricate interplay between the centromere, kinetochore, and spindle microtubules. Despite its importance, the architecture of this interface remains elusive. Here we used *in situ* cryo-electron tomography to visualize the native architecture of the kinetochore-microtubule interface in human U2OS cells at different stages of mitosis. We find that the centromere forms a pocket-like structure around kinetochore microtubules. Two morphologically distinct fibrillar densities form end-on and side-on connections to the plus-ends of microtubules within this centromeric pocket. Our data suggest a dynamic kinetochore-microtubule interface with multiple interactions between outer kinetochore components and spindle microtubules.

## Main

In mitosis, error-free segregation is mediated by a complex multi-protein assembly called the kinetochore, which connects the chromosome to spindle microtubules (MTs)^1–3^. Chromosome missegregation results in aneuploidy, which is associated with birth defects and cancers^4,5^. The structure of the kinetochore has been pursued for decades. In vertebrates, the kinetochore consists of more than a hundred proteins, and it binds 10-30 microtubules that make up the kinetochore-fiber (k-fiber)^6,7^. Classical electron microscopy (EM) studies of glutaraldehyde-fixed cells from diverse organisms revealed the kinetochore as a trilaminar structure comprising three morphological domains, including electron-dense inner and outer plates^8–10^. More recent EM studies using high-pressure freezing and freeze-substitution of PtK1 cells challenged this classic trilaminar view and instead found that the outer plate of the kinetochore is a network of long fibers oriented in the plane of the plate, rather than a dense disk^11^. Further work using similar techniques in PtK1 cells reported slender fibrils from the inner kinetochore attaching to the curved ends of MT protofilaments (PFs)^12–14^. However, different groups applying the same technique to the same cell line reached different conclusions, highlighting the limitations of sample preparation and imaging techniques used to date^15^ and underscoring the need for higher-resolution imaging of near-native samples to resolve these ambiguities.

To address these questions and to develop a more complete picture of the kinetochore-MT interface, we combined cryo-correlative light and electron microscopy (cryo-CLEM), cryo-focused ion beam milling (cryo-FIB milling), and cryo-electron tomography (cryo-ET) to visualize the native three-dimensional (3D) structure of the human kinetochore *in situ* in mitotic U2OS cells. We found that the centromere in U2OS cells has a pocket-like structure that accommodates multiple kinetochore microtubules (KMTs). This structure persisted throughout mitosis, and we observed sparsely-distributed nucleosome densities within the pocket. Our data reveals previously unresolved features of the centromere and kinetochore in mammalian cells, including two morphologically distinct fibrillar densities which form end-on and side-on connections to the plus-ends of microtubules within the centromeric pocket. Our findings suggesting that the pocket configuration scaffolds a dynamic kinetochore-microtubule interface in which multiple interactions facilitate stable attachment to MT plus-ends that are continually switching between growing and shrinking states.

## Results

### Workflow for cryo-electron tomography of kinetochores in mitotic U2OS cells

Cryo-ET has become the standard technique for visualizing the 3D organization of macromolecules in an unperturbed cellular context, achieved by cryopreserving samples in vitreous ice^16,17^. It has been successfully applied to study whole bacterial cells^18^, viruses^19^, and the thin edges of adherent eukaryotic cells^20^. However, the nuclear region of most mammalian cells exceeds 2 µm in thickness, precluding direct visualization by cryo-ET. Recently, cryo-FIB milling has been developed as a way to remove excess cellular material around a region of interest without the artifacts of cryo-ultramicrotomy^21–23^. We therefore sought to use cryo-FIB milling to expose kinetochores for cryo-ET. To locate cells at different mitotic stages, we grew U2OS cells on gold EM finder grids for 36 hours and added Hoechst stain shortly before plunge-freezing. The positions of mitotic cells were identified by cryo-fluorescence light microscopy (cryo-LM) and recorded relative to visible landmarks on the finder grid^24^ (Fig. 1A). Cryo-LM imaging allowed unambiguous identification of each cell’s mitotic stage (Fig. S1). Next, we located the same mitotic cells in the FIB/SEM instrument by aligning the fluorescence and SEM images using the grid bars and other markers (Fig. 1B). Since the limited axial resolution of cryo-LM imaging precluded us from determining the exact location of kinetochores along the z-axis, we cryo-FIB milled material above and below the center of the cell, leaving a thin (< 250 nm) lamella extending through the chromosome-containing region of the cell (Fig. 1C, D).

**Fig. 1.**
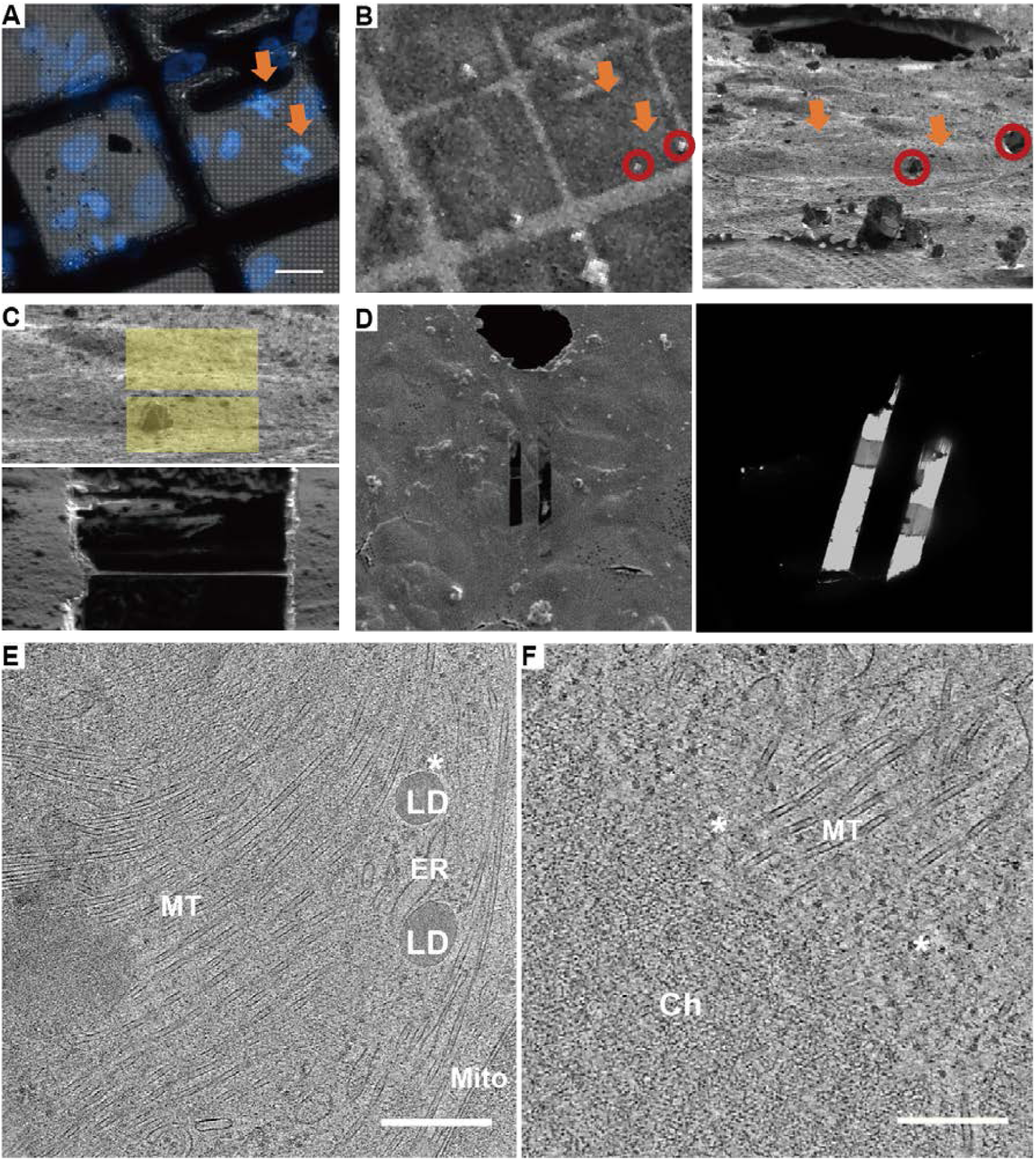
Experimental workflow developed to image mitotic U2OS cells by cryo-ET. **(A)** Cryo-light microscopy of U2OS cells grown on an EM grid. Orange arrows indicate mitotic cells. Scale bar: 30 µm. **(B)** SEM and FIB image of the same area on the grid as in **(A)**. Orange arrows point to the same mitotic cells as in (A). Red circles show the ice crystals on the grid that can be used for alignment. **(C)** A lamella through the central region of the cell made using cryo-FIB milling. Yellow areas indicate the milled regions. **(D)** SEM and TEM images of a milled region showing two lamellae. **(E)** Tomographic slice of a prometaphase cell thinned by cryo-FIB milling showing spindle microtubules. **(F)** Slice through a tomogram showing the mitotic chromosome and microtubules. MT: microtubules; Asterisks: ribosomes; Mito: mitochondrion; LD: lipid droplets; ER: endoplasmic reticulum; Ch: chromosome. Scale bars: 300 nm.

Finally, we imaged the lamellae by cryo-ET. The samples were well preserved, and we identified characteristic features of mitotic cells including ribosomes, spindle MTs, endoplasmic reticulum (ER), and condensed chromosomes (Fig. 1E, F, Movies S1, S2). We observed that the surface of the mitotic chromosomes was usually coated with a layer of ribosomes (Fig. 1F), consistent with previous reports^25,26^. The structure of condensed chromosomes and their relationship to spindle MTs were clearly visible, allowing us to distinguish between MTs that do not associate with kinetochores (non-KMTs) and those that do (KMTs) (Fig. 2). We collected hundreds of tomograms from cells in different mitotic stages and found 5 tomograms in which the plus-ends of KMTs were embedded in centromeric chromatin. These tomograms came from cells in prometaphase (n=1), metaphase (n=2), and anaphase (n=2). The complex molecular organization at the chromosome-microtubule interface is demonstrated in Fig. 2. The mitotic chromosome exhibits regions of dense material with ribosomes distributed on its surface following breakdown of the nuclear membrane. A bundle of kinetochore microtubules can be seen passing through fenestrations in the ER membrane and connecting with the mitotic chromosome at the centromere.

**Fig. 2.**
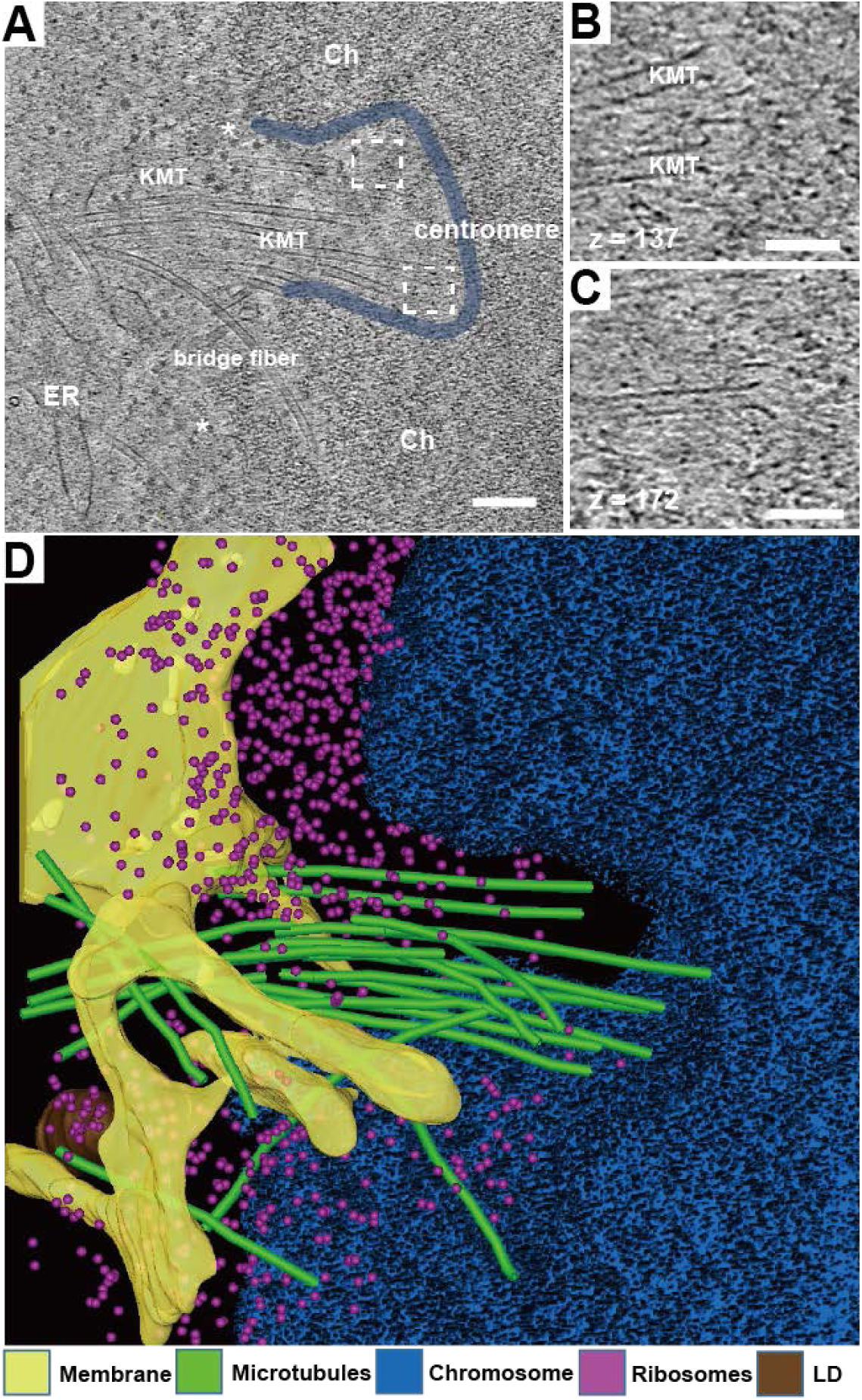
High-resolution architecture of the chromosome-microtubule interface in U2OS cells by cryo-ET. **(A)** Tomographic slice of a metaphase cell thinned by cryo-FIB milling showing an example kinetochore. Ch: chromosome; ER: endoplasmic reticulum; KMT: kinetochore microtubule; Asterisks: ribosomes. Scale bar: 150 nm. **(B) (C)** Enlarged views of areas in the same tomogram at different z slices showing the plus-ends of kinetochore microtubules. Scale bars: 60 nm. **(D)** 3D segmentation of the tomogram in panel **(A)** with color key shown at the bottom. LD: lipid droplet.

### Centromeric chromatin forms a pocket-like structure in the mitotic chromosome

The centromere is a specific region of the chromosome visually recognizable by a primary constriction^27^ and epigenetically defined by the histone H3 variant CENP-A^28–30^. We found that the centromeres in all our cryo-tomograms exhibited the same pocket-like appearance, from prometaphase through anaphase (Figs. 3A-C). This contrasts with classical EM studies of glutaraldehyde-fixed cells, where centromeres exhibit a variety of shapes, from pocket-like to plate-like to dome-shaped structures^12,15,31,32^. The width of the centromeric pocket in our tomograms ranged from 350-500 nm, and multiple KMTs were embedded in each pocket (Figs. S2, Movie S3). It has been estimated that mammalian kinetochores are attached to 10-30 MTs, collectively referred to as the k-fiber^33,34^. By digitally segmenting microtubules and the surface of the condensed chromosome in 3D, we found that the microtubules in a k-fiber are not always parallel and can exhibit different orientations (Figs. 2D, S2). Interestingly, some microtubules in the k-fiber did not connect to the chromosome but instead passed through it like non-KMTs (Figs. 3B, C, S2, yellow microtubules). They are likely to be the bridging fibers that connect sister kinetochore fibers as suggested by recent studies^35,36^. While the plus-ends of KMTs in the k-fiber could be found in different parts of the centromeric pocket, all KMTs with a visible plus-end in our tomograms were contained in the pocket from prometaphase to anaphase (Fig. S2).

**Fig. 3.**
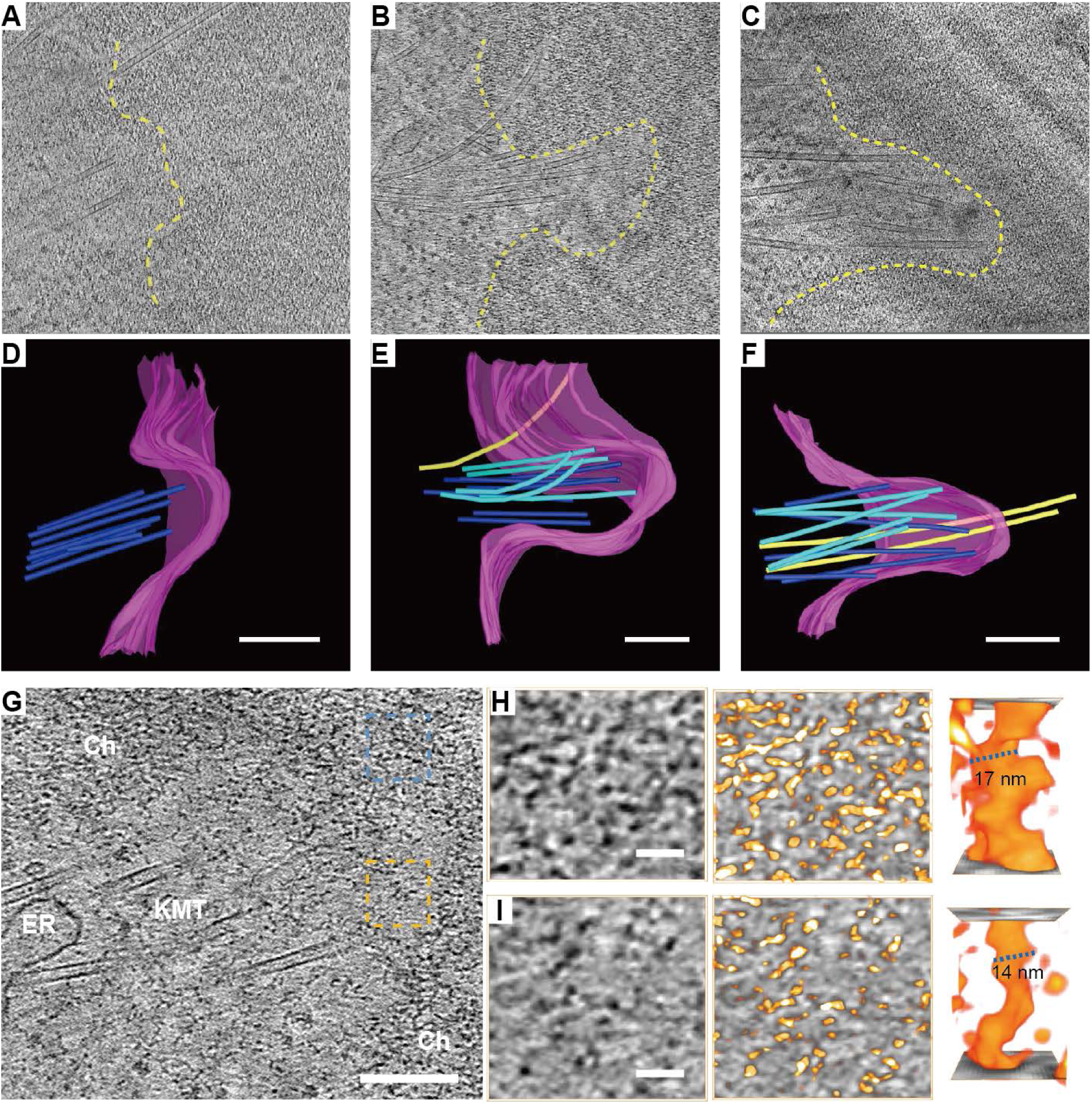
Centromeric chromatin forms a pocket-like structure in the mitotic chromosome. **(A, B, C)** Tomographic slices of cryo-FIB-milled U2OS cells at different mitotic stages containing kinetochores. **(D, E, F)** 3D segmentations of the corresponding tomograms. Magenta: chromosome surface; dark blue: KMTs with plus-ends visible in the tomogram; light blue: KMTs without visible plus-ends in the tomogram; yellow: bridging fibers. Scale bars: 300 nm. **(G)** Representative tomographic slice of a kinetochore with clear nucleosome densities. Ch: chromosome; ER: endoplasmic reticulum; KMT: kinetochore microtubule. Blue box: condensed chromosome; yellow box: centromeric pocket. Scale bar: 150 nm. **(H, I)** Left: magnified images of chromatin in the blue and yellow boxed areas, respectively, in **(G)**; middle: isosurface rendering of chromatin density in 3D; right: enlarged 3D views of a nucleosome chain. Scale bars: 30 nm.

Comparing the structure of centromeres at different mitotic stages, we found that metaphase and anaphase cells had a deeper centromeric pocket (600 nm, n=4) than prometaphase cells (200 nm, n=1) (Fig. 3A-C). Previous studies using fluorescence microscopy have shown that the kinetochore undergoes a rearrangement after the attachment of KMTs^37^. The shallow centromeric pocket we observed in prometaphase likely represents an early stage of formation of the primary constriction.

### Nucleosomes exhibit no higher-order structure at the centromere

A crucial open question about the structure of the centromere is how linear interspersion of CENP-A and H3 can be reconciled with the exclusion of H3 on chromosomes. It has been proposed that centromeric chromatin forms a unique higher-order structure, with CENP-A packaged either into an amphipathic-like solenoidal superhelix or as radial loops, with CENP-A nucleosomes clustered on the outer surface of the H3 chromatin^38,39^. However, details of the 3D organization of centromeric chromatin are still lacking. Our data challenges these models because we did not observe any higher-order structure of nucleosomes either at the centromere or in other parts of the chromosome, consistent with previous EM and X-ray scattering studies^40,41^.

The condensed chromatin in our tomograms appeared to be composed of disordered chains with a diameter of 10-20 nm (Fig. 3G, H). Interestingly, we found nucleosome densities in the centromeric pocket which were less condensed than in surrounding condensed chromatin, although they formed the same disordered 10-20 nm-wide chains as in the condensed chromosome (Fig. 3H, I). To determine whether this is a unique feature of the centromere, we compared nearby regions on the surface of non-centromeric chromatin, where we did not observe the same nucleosome chain structure (Fig. S3).

### Fibrils connect KMTs to chromatin through both side-on and end-on attachments

Next, we sought to find the structure that connects KMTs with the chromosome. The firm attachment between MTs and the kinetochore serves as the basis for chromosome movement. There are two major protein interaction networks in the kinetochore: the constitutive centromere-associated network (CCAN) and the Knl1 complex-Mis 12 complex-Ndc80 complex network (KMN)^42–44^. The Ndc80 complex of the KMN is central to kinetochore-MT attachment, directly binding to tubulin subunits of the MTs, along with other MT-binding proteins^45,46^. The Ndc80 complex is a heterotetramer of Spc25, Spc24, Nuf2, and Ndc80. Purified Ndc80 is a 57-nm-long rod with globular densities at either end^47,48^. Diverse models have been proposed for how the kinetochore connects KMTs to the chromosome. In the conformational wave model, curling PFs pull continuously on the kinetochore. Theoretical analysis suggests that this mechanism can produce sufficient force for movement^49^. Alternatively, the biased diffusion model posits that the kinetochore consists of multiple elements that form diffusive attachments to the microtubule^50^. The extent to which these theoretical models reflect the real architecture of the interface between MTs, and the kinetochore is unclear.

Analysis of our cryo-tomograms revealed two major modes of structural interaction between MTs and the kinetochore. In the first, we found fibril densities connected to the wall of MTs (Fig. 4A, C). This observation is consistent with *in vitro* experiments showing that several outer kinetochore components, including Ndc80, can directly bind to the wall of KMTs^51,52^. However, no such lateral connections of fibrils were seen *in situ* in previous work^14^. We also observed motor-protein-like densities on the walls of KMTs (Fig. S4), which were also unresolved in high-pressure-frozen samples. This highlights the power of cryo-ET for visualizing native cellular structure. To rule out the possibility that the side-on connections we observed were simply electron densities from the chromosome, we also examined non-KMTs passing through chromosomes. Interestingly, we saw that there is usually a translucent zone between the chromosome and the walls of non-KMTs, and no fibrillar densities were seen in this zone (Fig 4B, S5). This suggests that the side-on fibrils we observed are specific to the kinetochore.

**Fig. 4.**
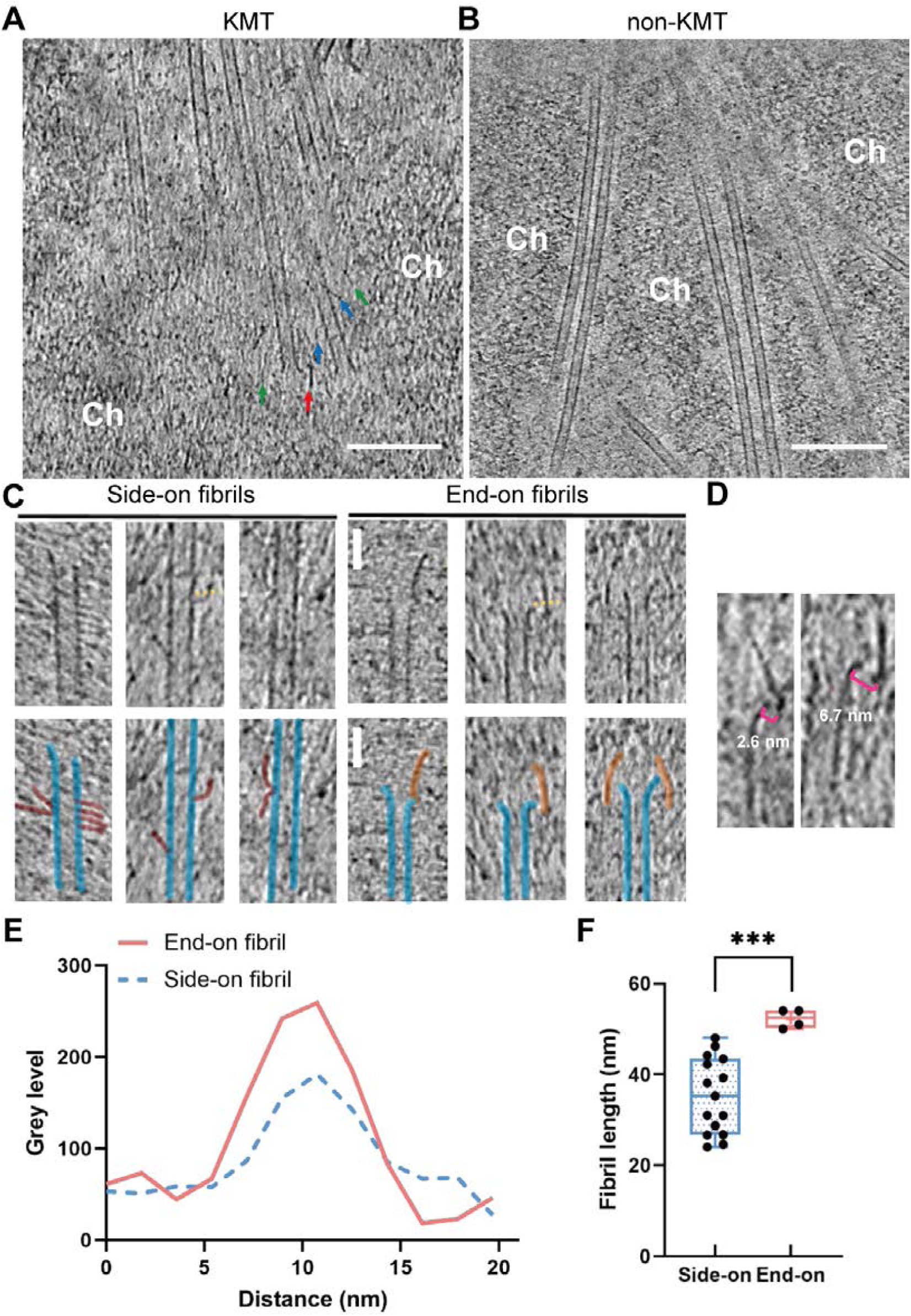
Fibrils connect KMT PFs to nearby chromatin through both end-on and side-on attachments. **(A)** Representative tomographic slice from a cryo-FIB-milled mitotic cell showing KMTs and the chromosome. Ch: chromosome; red arrowheads: end-on fibrils; blue arrowheads: side-on fibrils; green arrowheads: fibril densities not connected to microtubules. **(B)** Tomographic slice of non-KMTs passing through the mitotic chromosome region. Ch: chromosome. Scale bars: 100 nm. **(C)** Tomographic slices of KMTs. Side-on and end-on fibrils associated with PFs are indicated by graphic overlays below. Scale bars: 30 nm. **(D)** Typical distances between end-on fibrils and PFs. **(E)** Line density profile showing the thickness of end-on fibrils and side-on fibrils. **(F)** Boxplot representation of fibril length of side-on and end-on fibrils. Median values are marked by the central line. ***p<0.0001 by Student’s t-test.

The second mode of interaction we observed was mediated by fibrils near the flared plus-ends of KMTs, usually oriented parallel to the MT axis (Fig. 4A, C). A previous study suggested that lattice defects may cause PFs to separate from the MT wall^53^. To rule out the possibility that these end-on fibrils are remnants of disassembled MT PFs, we looked for lattice defects in the walls of both KMTs and non-KMTs. In both cases, lattice defects caused the microtubule to bend with an unusual curvature, but no fibril-like densities were seen (Fig. S6). The end-on fibrils we observed were thicker and more electron dense (darker in images) than side-on fibrils (Fig. 4E). The average length of end-on fibrils was 52 nm, which is consistent with the length of Ndc80 (57 nm). Side-on fibrils were less straight and more flexible in their shape and orientation (Fig. 4C), making it difficult to determine their length accurately, but they were shorter than end-on fibrils (Fig. 4F). Both classes of fibrils we observed–thick fibrils at the plus-end and thin fibrils on the wall of KMTs–localized exclusively to the centromeric region and thus are likely components of the outer kinetochore network.

### KMT PFs exhibit unique curvature with thick fibrils near their outer surface

Previous EM studies of mitotic cells suggested that there are fibrillar densities connected to the lumenal side of curved PFs. These fibrils appear to impede PF bending, suggesting a mechanism for converting the energy of MT depolymerization into chromosome movement^13,14^. Unlike previous reports, we did not observe fibrils binding to the lumenal side of flared KMT plus-ends. Instead, our cryo-tomograms showed the fibrils positioned close to the terminal tubulin unit of the curved PF, on the outer side (Fig. 4D). In our tomograms, we did not find any direct connections between the end-on fibrils and PFs. This is probably because the connection is very transient; once the tubulin unit dissociates from the PF, the attachment is lost. This observation resolves a key problem of the previous kinetochore fibril model: there was no obvious molecular candidate for these fibrils since both Ndc80 and CENP-E are known to bind the walls of MTs, not the lumen^51,54^.

Next, we examined the shapes of PFs from both KMTs and non-KMTs in our tomograms (Fig. S7). It is thought that the force for chromosome movement comes from MT dynamics, which are associated with GTP hydrolysis^55–58^. During growth, polymerizing MTs usually exhibit a straight form^59^. By contrast, during depolymerization, individual PF strands of tubulin dimers curve away from the central axis, flaring the MT plus-end^60^. It is believed that this morphological change is harnessed by outer kinetochore components, either a ring structure in yeast^61^ or a fibrillar structure in higher vertebrates^13^, to drive chromosome motion^42^. Both straight and curved forms of PFs can be found in MTs due to constant cycles of polymerization and depolymerization. Interestingly, we found that the PFs at the plus-ends of KMTs were always curved in the presence of thick end-on fibrils (n=5) (Fig. 4C), suggesting a relationship between those fibrils and MT depolymerization. Moreover, none of these PFs showed the long-curved form we saw in non-KMTs (Fig. S7C), indicating that the curling of these PFs is impeded by the fibrils. The fact that we did not observe thick fibrils near KMTs with a straight form suggests that the end-on fibrils may only associate with depolymerizing plus-ends.

## Discussion

In this study, we combined cryo-CLEM, cryo-FIB milling, and cryo-ET to visualize the unperturbed 3D ultrastructure of the kinetochore-microtubule interface in mitotic U2OS cells. Our method eliminates the harsh sample preparation steps of traditional EM and delivers the most native picture to date of this structure inside the cell. Our cryo-tomograms revealed a pocket structure of the centromere that accommodated multiple KMTs and deepened throughout mitosis, from prometaphase to anaphase. A pocket-shaped centromere could provide several advantages. First, it may increase the efficiency of MT capture by guiding MTs into the correct orientation for docking. Second, it provides a scaffold for the lateral interactions between KMTs and the kinetochore that we observed. Third, since MTs are undergoing constant assembly/disassembly during mitosis, the expanded interface of the pocket could prevent loss of disassembling KMTs from the kinetochore and facilitate their recapture.

Currently, resolution barriers hinder molecular identification of structural features seen in electron cryo-tomograms. Still, based on our results we can postulate a model, albeit speculative, for the molecular architecture of the vertebrate kinetochore. The thick fibrils we saw near the plus-ends of KMTs had a similar length (52 nm) and morphologically resembled the structure of Ndc80 (Fig. 4D). Recent *in vitro* studies have predicted that the Ndc80 complex form clusters, variously proposed to contain 4 or more than 10 subunits, on MTs, and only such clustered Ndc80 is thought to stably bind MTs and track shortening ends^46,51,62^. We therefore propose that the thick fibrils we saw are a form of clustered Ndc80. The side-on fibrils we observed were more flexible in morphology and orientation, thus making them unlikely to provide a rigid connection between the chromosome and the disassembling plus-ends of KMTs. It has been proposed that some kinetochores initially associate with the side of MTs to capture and orient MTs in the early stages of mitosis^52,58,63^. However, our data reveals that even after MTs have been captured, side-on fibrils persist into later stages of mitosis. We think it likely that at least some of these side-on attachments to MT walls also correspond to the Ndc80 complex. The fibrils exhibited similar orientations and distances from the MT wall as those observed for Ndc80/Nuf2 dimers binding to *in*-*vitro*-assembled MTs^51^. Such laterally attached Ndc80s fibrils could then cluster once a load-bearing signal is sensed, as previous studies suggested^64,65^. Our findings challenge previous static view of the kinetochore-microtubule interface and represent a key step toward a holistic structural understanding of how the kinetochore harnesses the power of microtubule dynamics to drive chromosome segregation in higher vertebrates.

## Acknowledgments

We thank Dr. Songye Chen and Dr. Andrey Malyutin for technical assistance with cryo-electron microscopy. We thank Dr. Catherine M. Oikonomou for her very helpful comments on the manuscript. Cryo-electron microscopy was performed in the Beckman Institute Resource Center for Transmission Electron Microscopy at Caltech.

## Author contributions

W.Z. and G.J.J. conceived of the project. W.Z. and G.J.J. designed the experiments. W.Z. performed and interpreted the experiments. W.Z. and G.J.J. wrote the paper.

## Data and materials availability

All data is available upon request. Tomograms will be made publicly available through the Caltech Electron Tomography Database at https://etdb.caltech.edu.

## Supplementary Materials

Materials and Methods

Figs. S1 to S7

Movies S1 to S4

## Supplementary Materials

### Materials and Methods

#### Cell culture

Osteosarcoma U2OS cells (ATCC, Manassas, VA) were cultured in McCoy’s 5A Medium (ATCC, Manassas, VA) supplemented with 10% tetracycline-free fetal bovine serum (FBS; Takara Bio), and 100 U/ml penicillin/streptomycin. Cell cultures were maintained in a humidified 37 °C incubator with 5% CO_2_.

#### TEM grid preparation

For cryo-CLEM and cryo-ET, U2OS cells were plated onto fibronectin-coated 200-mesh gold R2/2 London finder Quantifoil grids (Quantifoil Micro Tools). Grids were sterilized by UV irradiation for 30 min and immersed in culture medium in a CO_2_ incubator for 30 min. Cells were lysed in flasks using 0.05% trypsin-EDTA and seeded on 4-8 pretreated gold finder grids in 35-mm dishes. Cells were kept in an incubator for 30-36 hours. Before vitrification, cells were stained with Hoechst 33342 at a concentration of 1 µg/ml (Thermo Fisher).

For vitrification, all grids were blotted from the back side of the film using Whatman filter paper strips and immediately plunged into a liquid ethane/propane mixture using an FEI Vitrobot Mark IV (FEI Company, Eindhoven, Netherlands). The Vitrobot was set to 37 °C, 94% humidity, and 1 s drain time. The frozen grids were stored in sealed boxes in liquid nitrogen until further processing

#### Cryo-light microscopy

The EM cartridges were transferred into a cryo-FLM stage (FEI Cryostage) mounted on a Nikon Ti inverted microscope. The grids were imaged using a 60X extra-long-working-distance air-objective (Nikon CFI S Plan Fluor ELWD 60X NA 0.7 WD 2.62–1.8 mm). Images were recorded using the NIS Elements software from AutoQuant (Nikon Instruments Inc., Melville, NY). Following cryo–light microscopy imaging, EM cartridges containing frozen grids were stored in liquid nitrogen.

#### Cryo-FIB milling

Cryo–light microscopy of cells expressing ER-localized PIS-GFP guided cryo-FIB milling to regions of interest in the x-y plane. Plunge-frozen grids were then mounted in custom-modified Polara cartridges with channels milled through the bottom. This allowed samples to be milled at a low angle of incidence (∼10° to 12°) with respect to the carbon surface. These modified cartridges were transferred into an FEI Versa 3D equipped with a Quorum PP3010T Cryo-FIB/SEM preparation system (Quorum Technologies LLC, East Sussex, UK). The stage temperature was held at −186°C for all subsequent steps. Samples were sputter coated with a thin layer of platinum (15 mA, 60 s) before milling to minimize curtaining and to protect the front edge of the sample during milling. The average platinum thickness on the grid squares used in this study was approximately 50 nm, which agreed well with the set thickness in the sputter coater. Vitrified cells lying approximately perpendicular to the focused ion beam were located via SEM. The vitrified samples were imaged at 5 to 10 keV with the SEM and milled with 30-keV gallium ions by scanning the regions of interest. A 0.3 nA beam current was used for rough milling. The current was stepped down slowly from 0.3 nA to 30 pA as the thickness of the lamellae decreased to 500 nm. In the end, 10 pA current was used to polish the lamellae and reduce the thickness of lamellae to under 300 nm.

#### Cryo-ET and tomogram reconstruction

Cryo-EM grids previously imaged by cryo-LM were subsequently imaged by cryo-ET using an FEI G2 Polara 300 keV FEG transmission electron microscope equipped with an energy filter (slit width 20 eV for higher magnifications: Gatan), and a 4 k × 4 k K2 Summit direct detector (Gatan) in counting mode. Cryo-tomograms were also collected using a Titan Krios transmission electron microscope (Thermo Fisher) equipped with a 300 keV field emission gun, energy filter (Gatan), and K2 or K3 Summit direct electron detector (Gatan).

Lamellae were located by making maps of the entire grid using the SerialEM software. Tilt-series acquisition was done with SerialEM with a 2° tilt increment for a total range of ±60° or ±50° (in two halves, separated at 0°). Individual tilt series were recorded at a magnification of ×19500 and a defocus of −6 to −10 μm. The cumulative dose of each tilt series was between 120 and 160 e^−^/Å^2^.

Tomograms were reconstructed using the IMOD software (http://bio3d.colorado.edu/imod/). In the absence of fiducial gold nanoparticles in the FIB-milled lamellae, alignment of tilt-series projection images was performed with patch-tracking. Final alignment of the tilt-series images was performed using the linear interpolation option in IMOD without contrast transfer function (CTF) correction. Aligned stacks were low-pass filtered (0.35, σ = 0.05) to eliminate high-frequency noise. Weighted back-projection reconstruction was performed, and the SIRT-like filter was used with 8 iterations.

#### Tomogram visualization and segmentation

Cryo-tomograms were subsequently segmented using the Amira software package (Thermo Fisher Scientific, FEI). Segmentation was performed manually using density thresholds. Morphological measurements of segmented data were also performed in Amira.

#### Statistical Analysis

Statistical analyses were performed with Fiji and Prism (GraphPad). The length of fibrils was assessed visually by tracing the density in 3D using Fiji. Given the subtlety of these structural features, the measurements should not be taken as precise. PFs were traced in IMOD, and points extracted using the *howflared* function in IMOD.

**Fig. S1.**
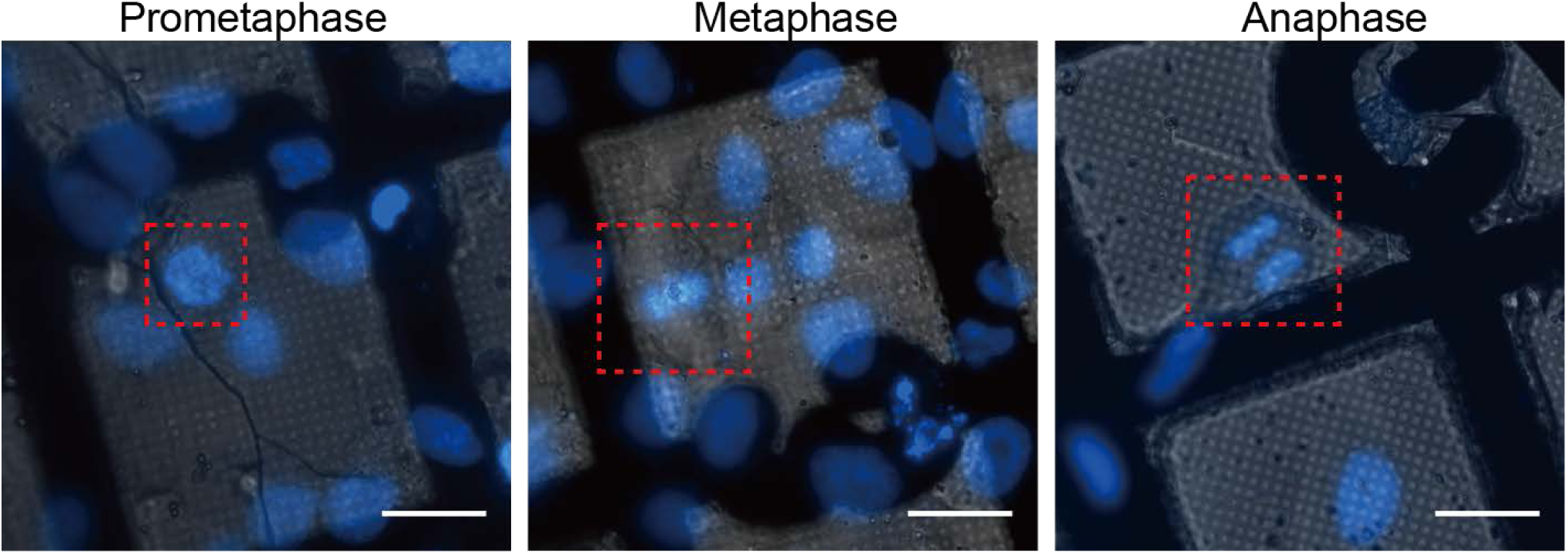
Cryo-LM of U2OS cells grown on EM grids captured at different mitotic stages. Scale bars: 30 µm.

**Fig. S2.**
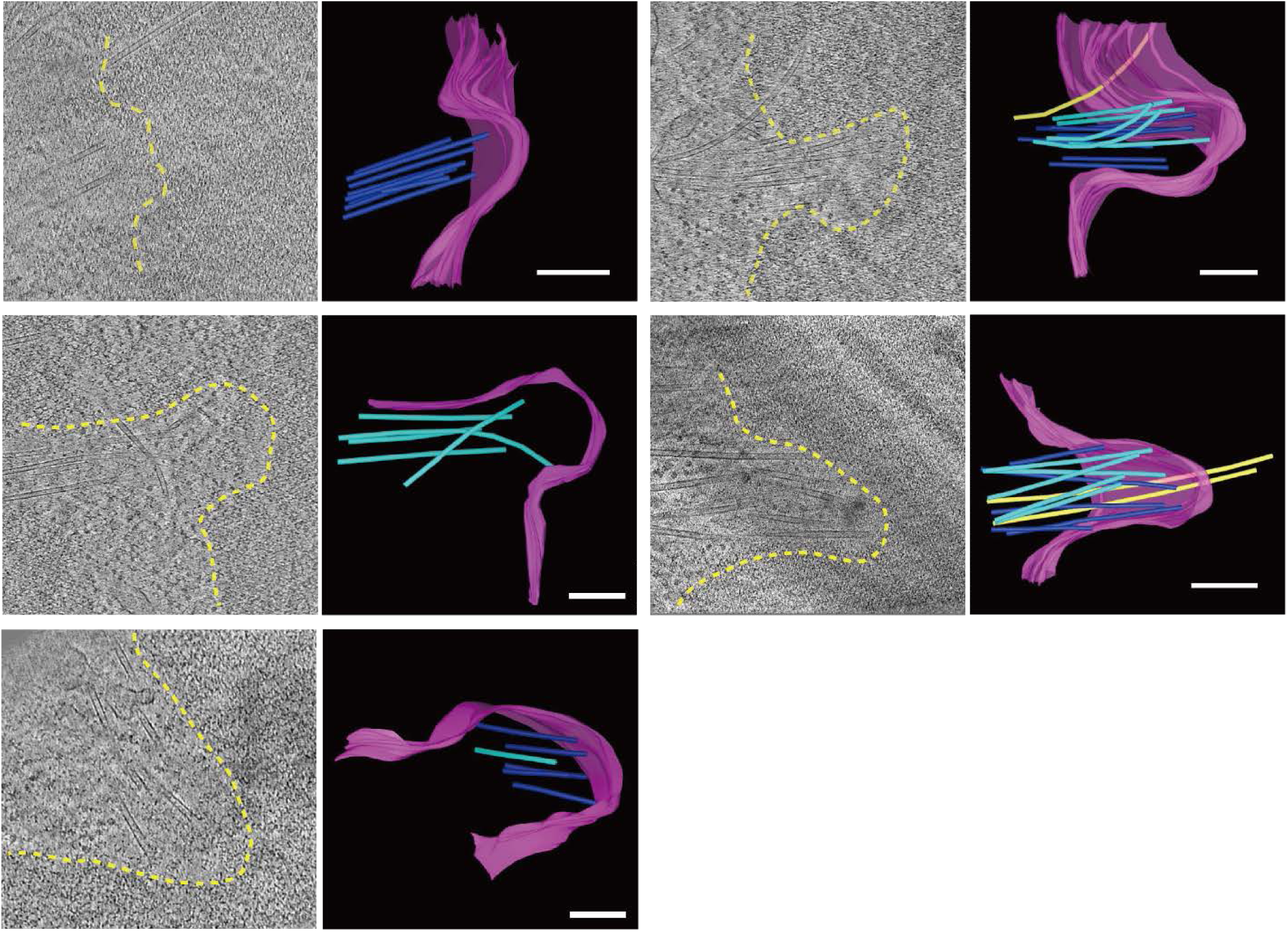
Gallery of segmentations of tomograms containing chromosome-microtubule interfaces. Yellow dashed line: trajectory of chromosome surface in the tomographic slice; magenta: chromosome surface; dark blue: KMTs with plus-ends visible in the tomogram; light blue: KMTs without visible plus-ends in the tomogram; yellow: bridging fibers. Scale bars: 300 nm.

**Fig. S3.**
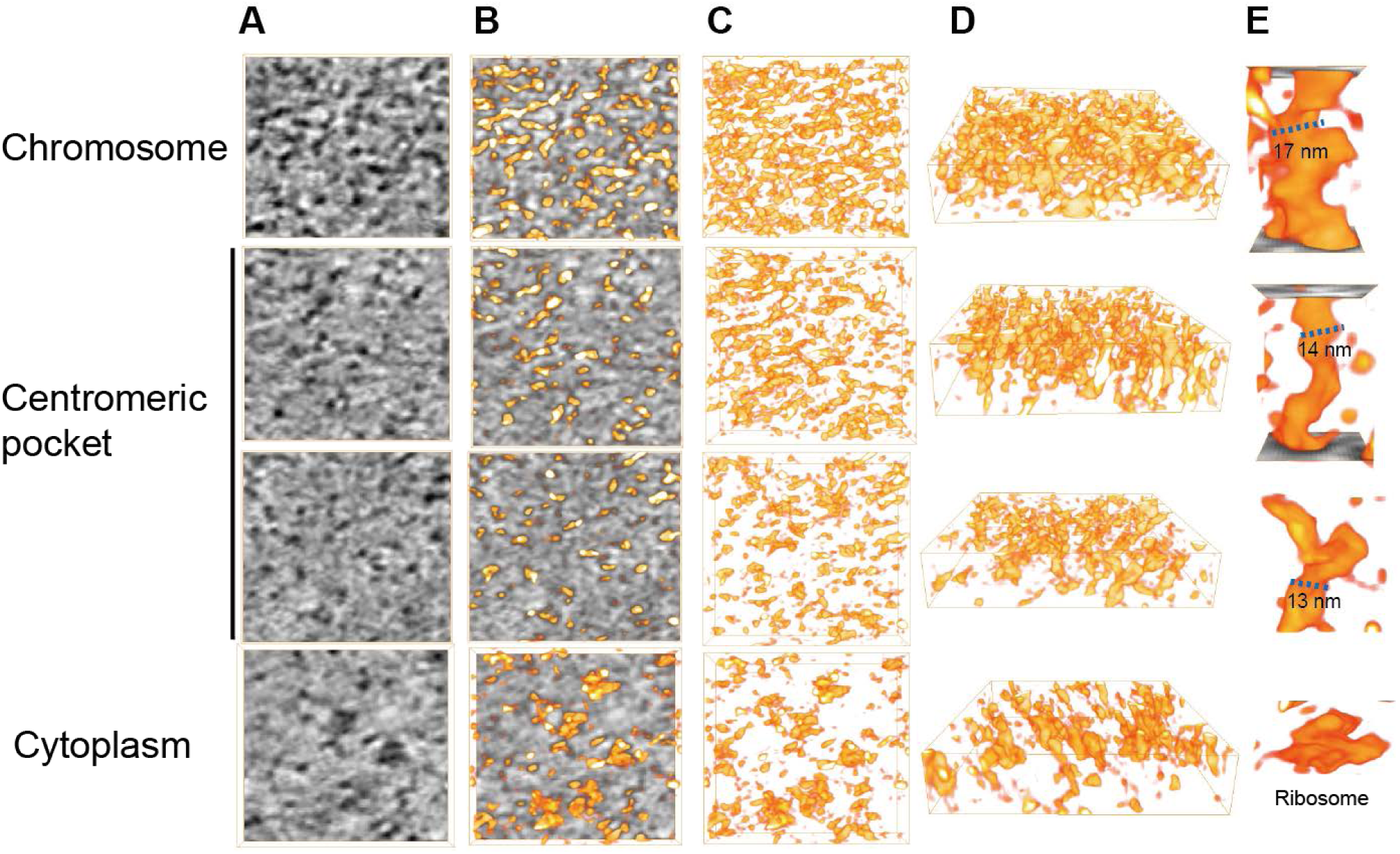
Chromatin is less condensed in the centromeric pocket than in neighboring chromosomal regions. **(A)** Tomographic slices showing densities in the indicated regions. Isosurface renderings of the densities are overlaid in **(B)** and isolated in **(C). (D)** Rotated views of the same surface renderings in **(C). (E)** Enlarged views of densities showing characteristic widths of nucleosome chains in condensed chromatin and the centromere pocket. Those nucleosome chains are not observed in the cytoplasm (bottom).

**Fig. S4.**
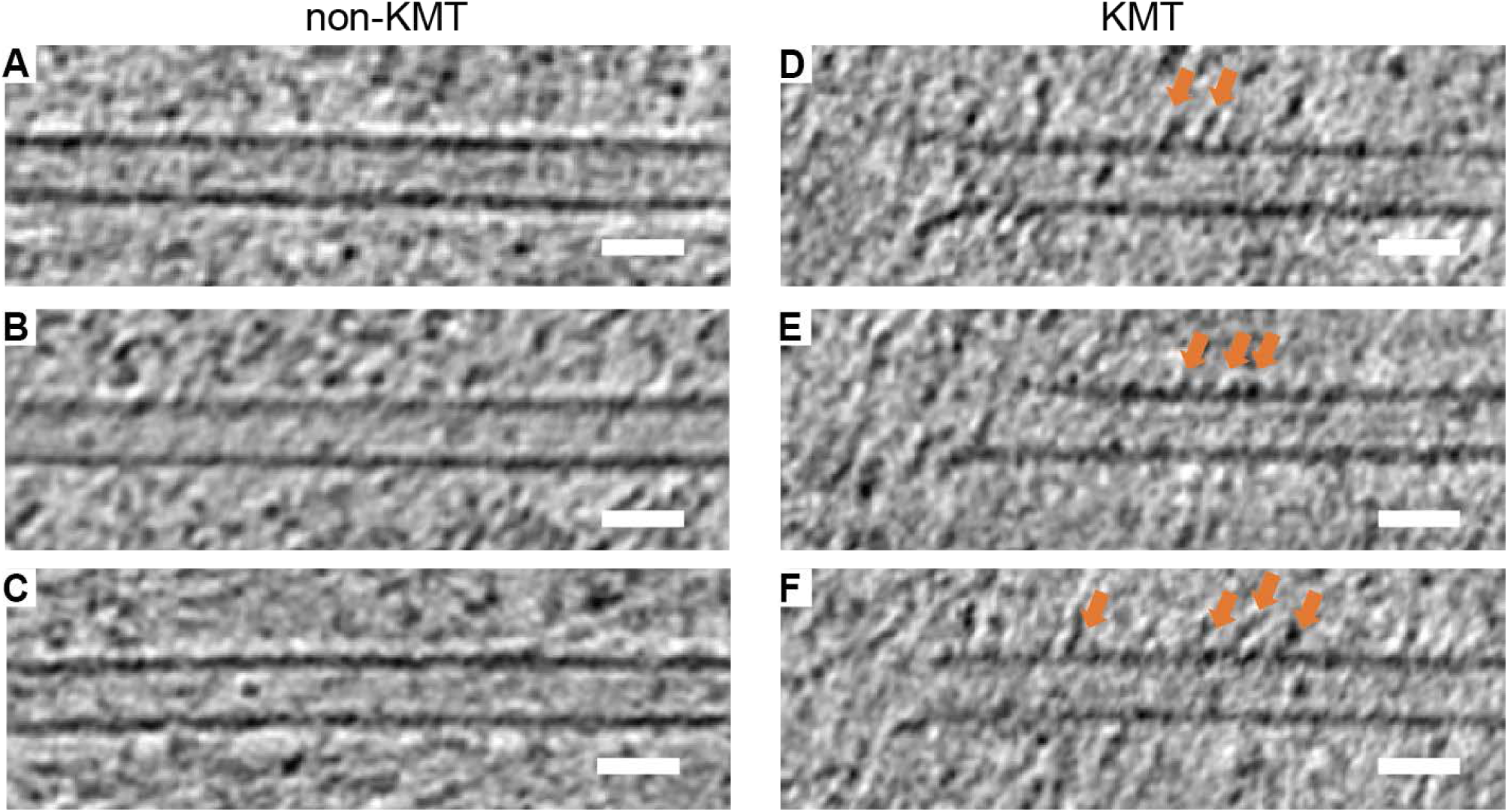
Motor-protein-like density on the walls of KMTs. (**A, B, C**) Representative tomographic slices of non-KMTs from mitotic U2OS cells thinned by cryo-FIB milling. (**D, E, F**) Tomographic slices of KMTs with motor-protein-like densities on their walls (orange arrows). Scale bars: 25 nm.

**Fig. S5.**
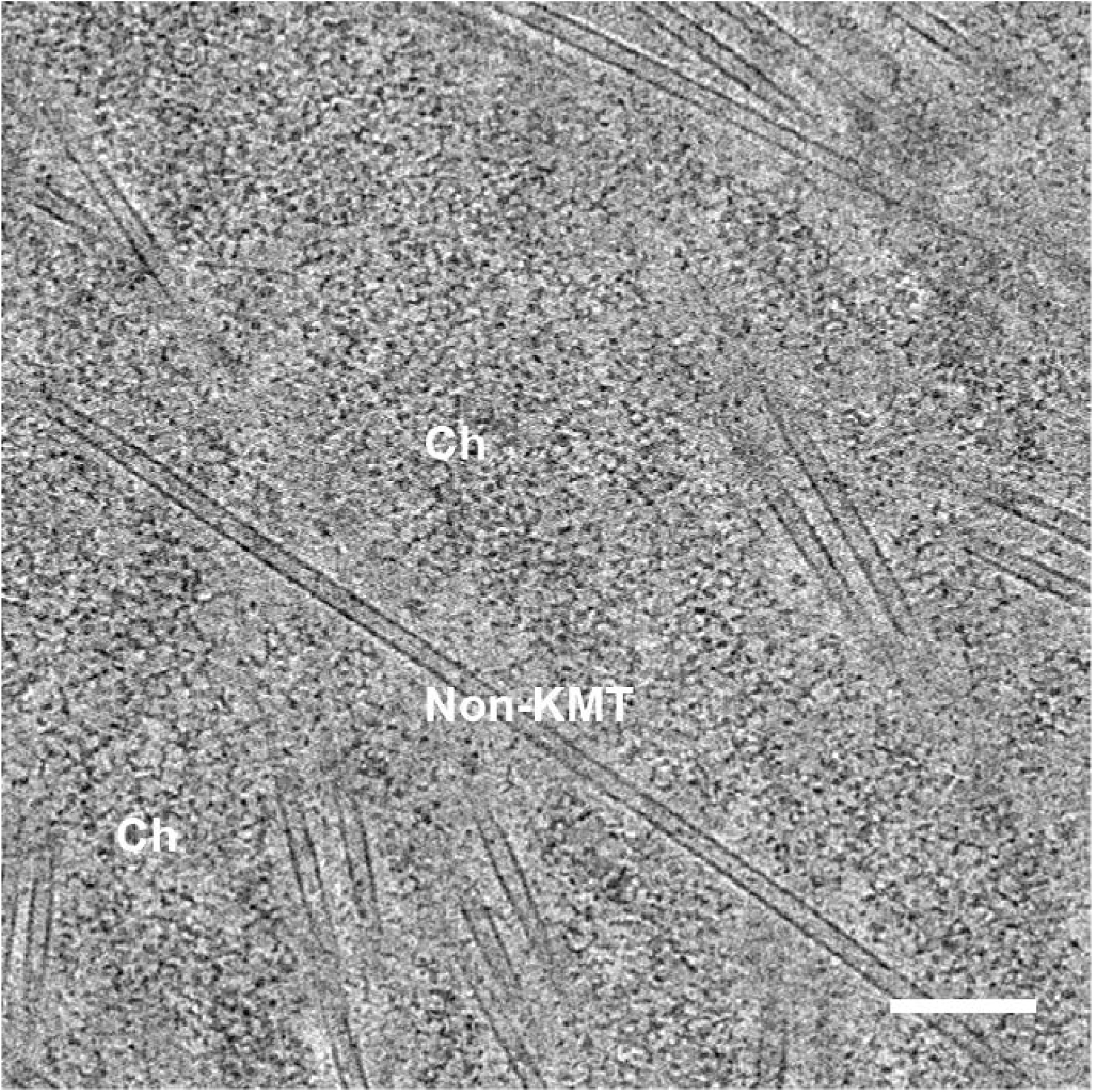
Non-KMT passing through the chromosome without any associated fibril densities. Scale bar: 150 nm.

**Fig. S6.**
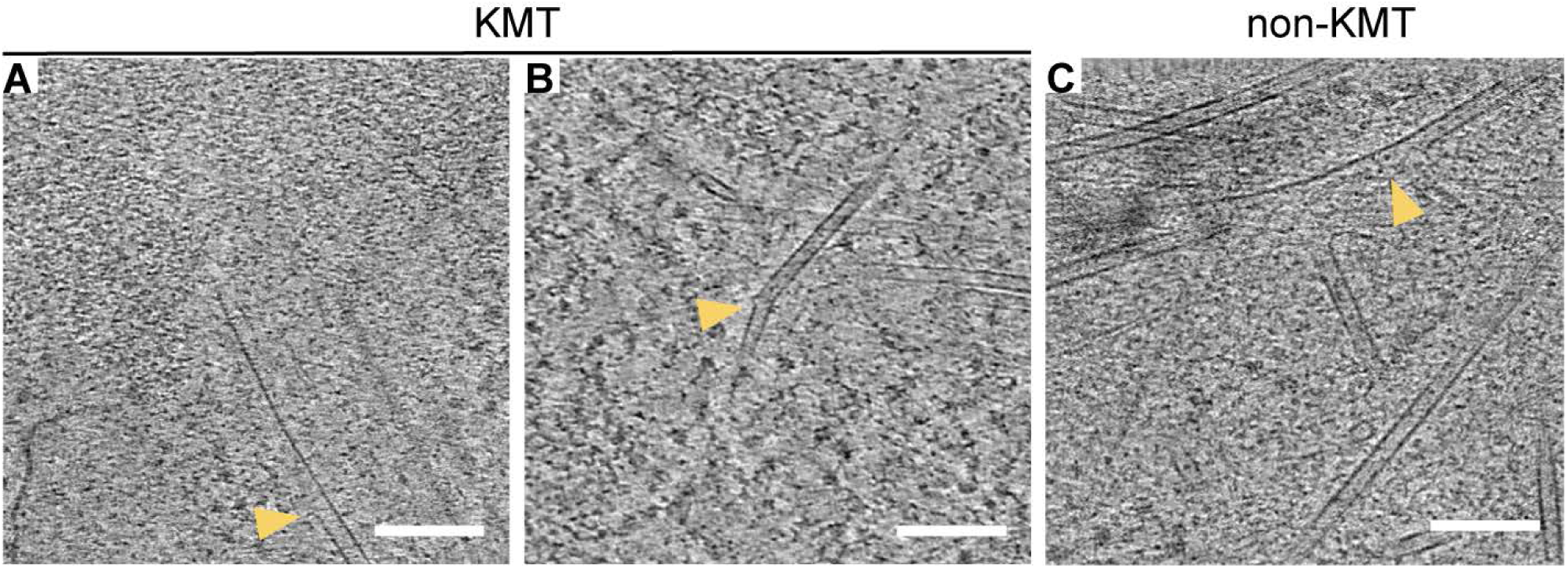
Microtubule lattice defects in both KMTs and non-KMTs observed by cryo-ET. Representative tomographic slices demonstrating lattice defects in KMTs **(A, B)** and non-KMTs **(C)** in mitotic U2OS cells. Arrowheads indicate defects caused by loss of tubulins from the MT wall. Scale Bar: 100 nm.

**Fig. S7.**
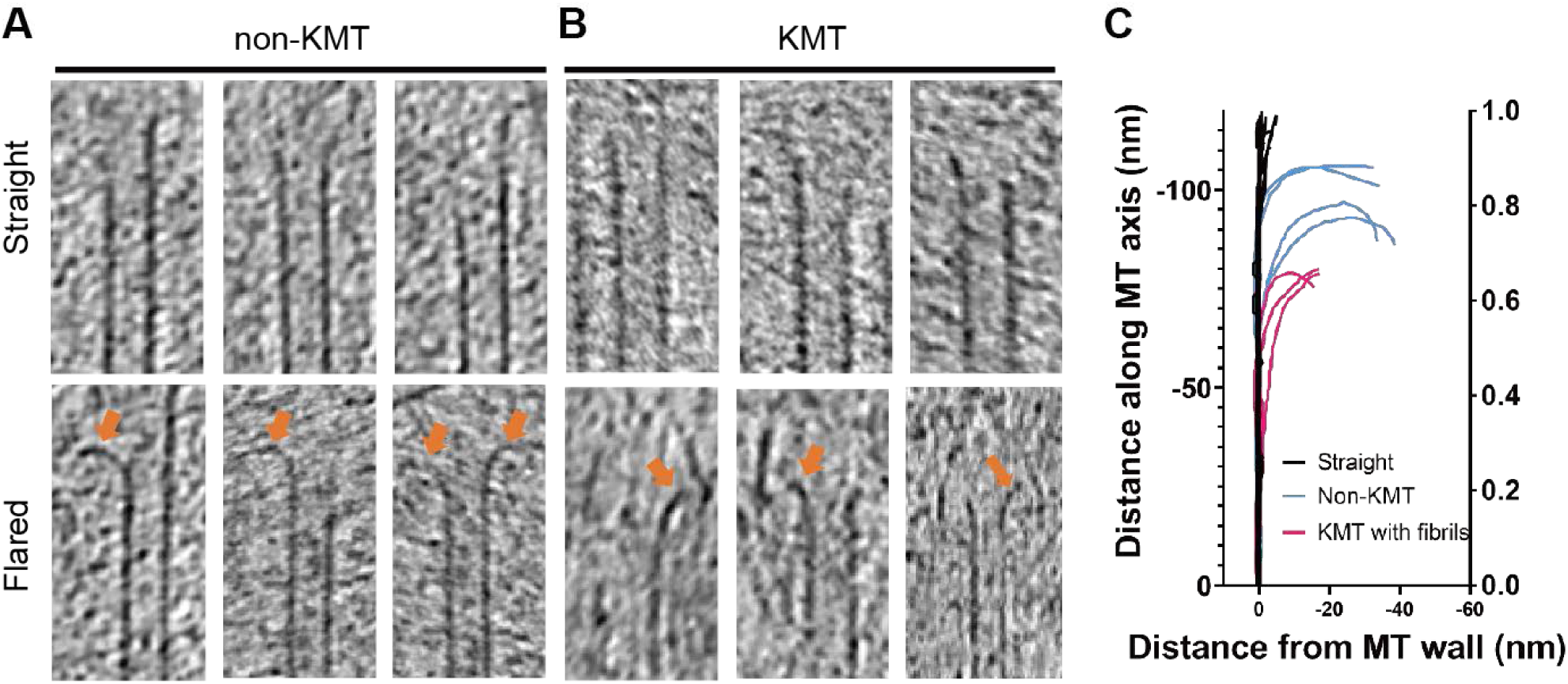
Comparison of flared plus-ends of KMTs and non-KMTs. Tomographic slices of plus-ends of non-KMTs **(A)** and KMTs **(B)** from mitotic U2OS cells. Orange arrows indicate flared PFs. **(C)** Shapes of PFs from MTs shown in (A, B). KMTs associated with end-on fibrils (magenta) have an intermediate curvature compared to flared non-KMTs (blue) and straight forms (black).

### Movies S1 to S4

**Movie S1. Tomogram of the prometaphase U2OS cell shown in Fig. 1E**.

The video scrolls up and down in Z through the tomographic volume. Scale bar: 300 nm.

**Movie S2. Tomogram of the condensed chromosome and nearby microtubules shown in Fig. 1F**.

The video scrolls up and down in Z through the tomographic volume. Scale bar: 300 nm.

**Movie S3. Tomographic volume of the kinetochore shown in Fig. 2**.

The video scrolls up and down in Z through the tomographic volume. Scale bar: 300 nm

**Movie S4. Segmentation of the kinetochore shown in Fig. 2**.

The video scrolls up and down in Z through the tomographic volume and reveals the 3D segmentation of identified structures. ER membrane: yellow; microtubules: green; chromosome: blue; ribosomes: pink; lipid droplet (LD): brown.

## References

1. McIntosh, J. R., Grishchuk, E. L. & West, R. R. Chromosome-Microtubule Interactions During Mitosis. Annual Review of Cell and Developmental Biology 18, 193–219 (2002).

2. Rieder, C. L. & Salmon, E. D. The vertebrate cell kinetochore and its roles during mitosis. Trends Cell Biol 8, 310–318 (1998).

3. Desai, A. & Mitchison, T. J. Microtubule polymerization dynamics. Annu Rev Cell Dev Biol 13, 83–117 (1997).

4. Pfau, S. J. & Amon, A. Chromosomal instability and aneuploidy in cancer: from yeast to man. EMBO Rep 13, 515–527 (2012).

5. Giam, M. & Rancati, G. Aneuploidy and chromosomal instability in cancer: a jackpot to chaos. Cell Div 10, (2015).

6. Cheeseman, I. M. The kinetochore. Cold Spring Harb Perspect Biol 6, a015826 (2014).

7. Pesenti, M. E., Weir, J. R. & Musacchio, A. Progress in the structural and functional characterization of kinetochores. Curr Opin Struct Biol 37, 152–163 (2016).

8. Rieder, C. L. The formation, structure, and composition of the mammalian kinetochore and kinetochore fiber. Int Rev Cytol 79, 1–58 (1982).

9. McEwen, B. F., Arena, J. T., Frank, J. & Rieder, C. L. Structure of the colcemid-treated PtK1 kinetochore outer plate as determined by high voltage electron microscopic tomography. J Cell Biol 120, 301–312 (1993).

10. Cooke, C. A., Schaar, B., Yen, T. J. & Earnshaw, W. C. Localization of CENP-E in the fibrous corona and outer plate of mammalian kinetochores from prometaphase through anaphase. Chromosoma 106, 446–455 (1997).

11. Dong, Y., Vanden Beldt, K. J., Meng, X., Khodjakov, A. & McEwen, B. F. The outer plate in vertebrate kinetochores is a flexible network with multiple microtubule interactions. Nature Cell Biology 9, 516–522 (2007).

12. McEwen, B. F., Hsieh, C. E., Mattheyses, A. L. & Rieder, C. L. A new look at kinetochore structure in vertebrate somatic cells using high-pressure freezing and freeze substitution. Chromosoma 107, 366–375 (1998).

13. McIntosh, J. R. et al. Fibrils Connect Microtubule Tips with Kinetochores: A Mechanism to Couple Tubulin Dynamics to Chromosome Motion. Cell 135, 322–333 (2008).

14. McIntosh, J. R. et al. Conserved and divergent features of kinetochores and spindle microtubule ends from five species. J Cell Biol 200, 459–474 (2013).

15. McEwen, B. F. & Dong, Y. Contrasting models for kinetochore microtubule attachment in mammalian cells. Cell. Mol. Life Sci. 67, 2163–2172 (2010).

16. Dobro, M. J., Melanson, L. A., Jensen, G. J. & McDowall, A. W. Plunge Freezing for Electron Cryomicroscopy. 481, 63–82 (2010).

17. Tao, C.-L. et al. Differentiation and Characterization of Excitatory and Inhibitory Synapses by Cryo-electron Tomography and Correlative Microscopy. J. Neurosci. 38, 1493–1510 (2018).

18. Oikonomou, C. M., Chang, Y.-W. & Jensen, G. J. A new view into prokaryotic cell biology from electron cryotomography. Nat Rev Microbiol 14, 205–220 (2016).

19. Luque, D. & Castón, J. R. Cryo-electron microscopy for the study of virus assembly. Nature Chemical Biology 16, 231–239 (2020).

20. Visualizing insulin vesicle neighborhoods in β cells by cryo–electron tomography | Science Advances. https://advances.sciencemag.org/content/6/50/eabc8258.

21. Rigort, A. & Plitzko, J. M. Cryo-focused-ion-beam applications in structural biology. Archives of Biochemistry and Biophysics 581, 122–130 (2015).

22. Marko, M., Hsieh, C., Schalek, R., Frank, J. & Mannella, C. Focused-ion-beam thinning of frozen-hydrated biological specimens for cryo-electron microscopy. Nature Methods 4, 215–217 (2007).

23. Rigort, A. et al. Focused ion beam micromachining of eukaryotic cells for cryoelectron tomography. Proceedings of the National Academy of Sciences 109, 4449–4454 (2012).

24. Chang, Y. W. et al. Correlated cryogenic photoactivated localization microscopy and cryo-electron tomography. Nature methods 11, 737–9 (2014).

25. Nishino, Y. et al. Human mitotic chromosomes consist predominantly of irregularly folded nucleosome fibres without a 30-nm chromatin structure. EMBO J 31, 1644–53 (2012).

26. Joti, Y. et al. Chromosomes without a 30-nm chromatin fiber. Nucleus 3, 404–410 (2012).

27. Fukagawa, T. & Earnshaw, W. C. The centromere: chromatin foundation for the kinetochore machinery. Dev Cell 30, 496–508 (2014).

28. Marshall, O. J., Marshall, A. T. & Choo, K. H. A. Three-dimensional localization of CENP-A suggests a complex higher order structure of centromeric chromatin. J Cell Biol 183, 1193–1202 (2008).

29. Foltz, D. R. et al. Centromere-specific assembly of CENP-a nucleosomes is mediated by HJURP. Cell 137, 472–484 (2009).

30. Okada, M., Okawa, K., Isobe, T. & Fukagawa, T. CENP-H-containing complex facilitates centromere deposition of CENP-A in cooperation with FACT and CHD1. Mol Biol Cell 20, 3986–3995 (2009).

31. Kixmoeller, K., Allu, P. K. & Black, B. The centromere comes into focus: from CENP-A nucleosomes to kinetochore connections with the spindle. Open Biology (2020) doi:10.1098/rsob.200051.

32. Brinkley, B. R. & Stubblefield, E. The fine structure of the kinetochore of a mammalian cell in vitro. Chromosoma 19, 28–43 (1966).

33. Walczak, C. E., Cai, S. & Khodjakov, A. Mechanisms of chromosome behaviour during mitosis. Nat Rev Mol Cell Biol 11, 91–102 (2010).

34. Chan, G. K., Liu, S.-T. & Yen, T. J. Kinetochore structure and function. Trends in Cell Biology 15, 589–598 (2005).

35. Vukušić, K. et al. Microtubule Sliding within the Bridging Fiber Pushes Kinetochore Fibers Apart to Segregate Chromosomes. Dev. Cell 43, 11–23.e6 (2017).

36. Kajtez, J. et al. Overlap microtubules link sister k-fibres and balance the forces on bi-oriented kinetochores. Nature Communications 7, 10298 (2016).

37. Renda, F. et al. kSHREC ‘Delta’ reflects the shape of kinetochore rather than intrakinetochore tension. BioRxiv 811075 (2019) doi:10.1101/811075.

38. Blower, M. D., Sullivan, B. A. & Karpen, G. H. Conserved Organization of Centromeric Chromatin in Flies and Humans. Dev Cell 2, 319–330 (2002).

39. Ribeiro, S. A. et al. A super-resolution map of the vertebrate kinetochore. PNAS 107, 10484–10489 (2010).

40. Ou, H. D. et al. ChromEMT: Visualizing 3D chromatin structure and compaction in interphase and mitotic cells. Science 357, eaag0025 (2017).

41. Maeshima, K., Imai, R., Tamura, S. & Nozaki, T. Chromatin as dynamic 10-nm fibers. Chromosoma 123, 225–237 (2014).

42. Musacchio, A. & Desai, A. A Molecular View of Kinetochore Assembly and Function. Biology 6, 5 (2017).

43. Hara, M. & Fukagawa, T. Critical Foundation of the Kinetochore: The Constitutive Centromere-Associated Network (CCAN). In Centromeres and Kinetochores: Discovering the Molecular Mechanisms Underlying Chromosome Inheritance (ed. Black, B. E.) 29–57 (Springer International Publishing, 2017). doi:10.1007/978-3-319-58592-5_2.

44. Varma, D. & Salmon, E. D. The KMN protein network – chief conductors of the kinetochore orchestra. J Cell Sci 125, 5927–5936 (2012).

45. Guimaraes, G. J., Dong, Y., McEwen, B. F. & Deluca, J. G. Kinetochore-microtubule attachment relies on the disordered N-terminal tail domain of Hec1. Curr Biol 18, 1778–1784 (2008).

46. Powers, A. F. et al. The Ndc80 Kinetochore Complex Forms Load-Bearing Attachments to Dynamic Microtubule Tips via Biased Diffusion. Cell 136, 865–875 (2009).

47. Wei, R. R., Sorger, P. K. & Harrison, S. C. Molecular organization of the Ndc80 complex, an essential kinetochore component. PNAS 102, 5363–5367 (2005).

48. Ciferri, C. et al. Architecture of the human ndc80-hec1 complex, a critical constituent of the outer kinetochore. J Biol Chem 280, 29088–29095 (2005).

49. Asbury, C. L., Tien, J. F. & Davis, T. N. Kinetochores’ Gripping Feat: Conformational Wave or Biased Diffusion? Trends Cell Biol 21, 38–46 (2011).

50. Hill, T. L. Theoretical problems related to the attachment of microtubules to kinetochores. Proc Natl Acad Sci U S A 82, 4404–4408 (1985).

51. Alushin, G. M. et al. The Ndc80 kinetochore complex forms oligomeric arrays along microtubules. Nature 467, 805–810 (2010).

52. Gonen, S. et al. The structure of purified kinetochores reveals multiple microtubule attachment sites. Nat Struct Mol Biol 19, 925–929 (2012).

53. Chakraborty, S., Mahamid, J. & Baumeister, W. Cryoelectron Tomography Reveals Nanoscale Organization of the Cytoskeleton and Its Relation to Microtubule Curvature Inside Cells. Structure 28, 991–1003.e4 (2020).

54. Wilson-Kubalek, E. M., Cheeseman, I. M., Yoshioka, C., Desai, A. & Milligan, R. A. Orientation and structure of the Ndc80 complex on the microtubule lattice. J. Cell Biol. 182, 1055–1061 (2008).

55. Coue, M., Lombillo, V. A. & McIntosh, J. R. Microtubule depolymerization promotes particle and chromosome movement in vitro. J Cell Biol 112, 1165–1175 (1991).

56. Koshland, D. E., Mitchison, T. J. & Kirschner, M. W. Polewards chromosome movement driven by microtubule depolymerization in vitro. Nature 331, 499–504 (1988).

57. Grishchuk, E. L. & McIntosh, J. R. Microtubule depolymerization can drive poleward chromosome motion in fission yeast. EMBO J 25, 4888–4896 (2006).

58. Tanaka, K., Kitamura, E., Kitamura, Y. & Tanaka, T. U. Molecular mechanisms of microtubule-dependent kinetochore transport toward spindle poles. J Cell Biol 178, 269–281 (2007).

59. Müller-Reichert, T., Chrétien, D., Severin, F. & Hyman, A. A. Structural changes at microtubule ends accompanying GTP hydrolysis: Information from a slowly hydrolyzable analogue of GTP, guanylyl (α,β)methylenediphosphonate. PNAS 95, 3661–3666 (1998).

60. Mandelkow, E. M., Mandelkow, E. & Milligan, R. A. Microtubule dynamics and microtubule caps: a time-resolved cryo-electron microscopy study. J Cell Biol 114, 977–991 (1991).

61. Jenni, S. & Harrison, S. C. Structure of the DASH/Dam1 complex shows its role at the yeast kinetochore-microtubule interface. Science 360, 552–558 (2018).

62. Ciferri, C. et al. Implications for kinetochore-microtubule attachment from the structure of an engineered Ndc80 complex. Cell 133, 427–439 (2008).

63. Kapoor, T. M. et al. Chromosomes Can Congress to the Metaphase Plate Before Biorientation. Science 311, 388–391 (2006).

64. Helgeson, L. A. et al. Human Ska complex and Ndc80 complex interact to form a load-bearing assembly that strengthens kinetochore-microtubule attachments. Proc. Natl. Acad. Sci. U.S.A. 115, 2740–2745 (2018).

65. Akiyoshi, B. et al. Tension directly stabilizes reconstituted kinetochore-microtubule attachments. Nature 468, 576–579 (2010).

